## Supplementary File for "Persistent Extrasynaptic Hyperdopaminergia in the mouse Hippocampus Induces Plasticity and Recognition Memory Deficits Reversed by Antipsychotics"

**Figure Sup 1:** Dopamine transporter expression in DA fibers to the Hippocampus

**Figure Sup 2:** Effect of GBR12935 intrahippocampal chronic infusion in the septal/dorsal HP on spatial and recognition memory.

**Supplemental Methods:**

- **Stereotaxic surgeries**
- **Immunohistolabeling and microscopy**
- **Intrahippocampal infusions**
- **Pharmacology**
- **Behavior**
- **Electrophysiology**

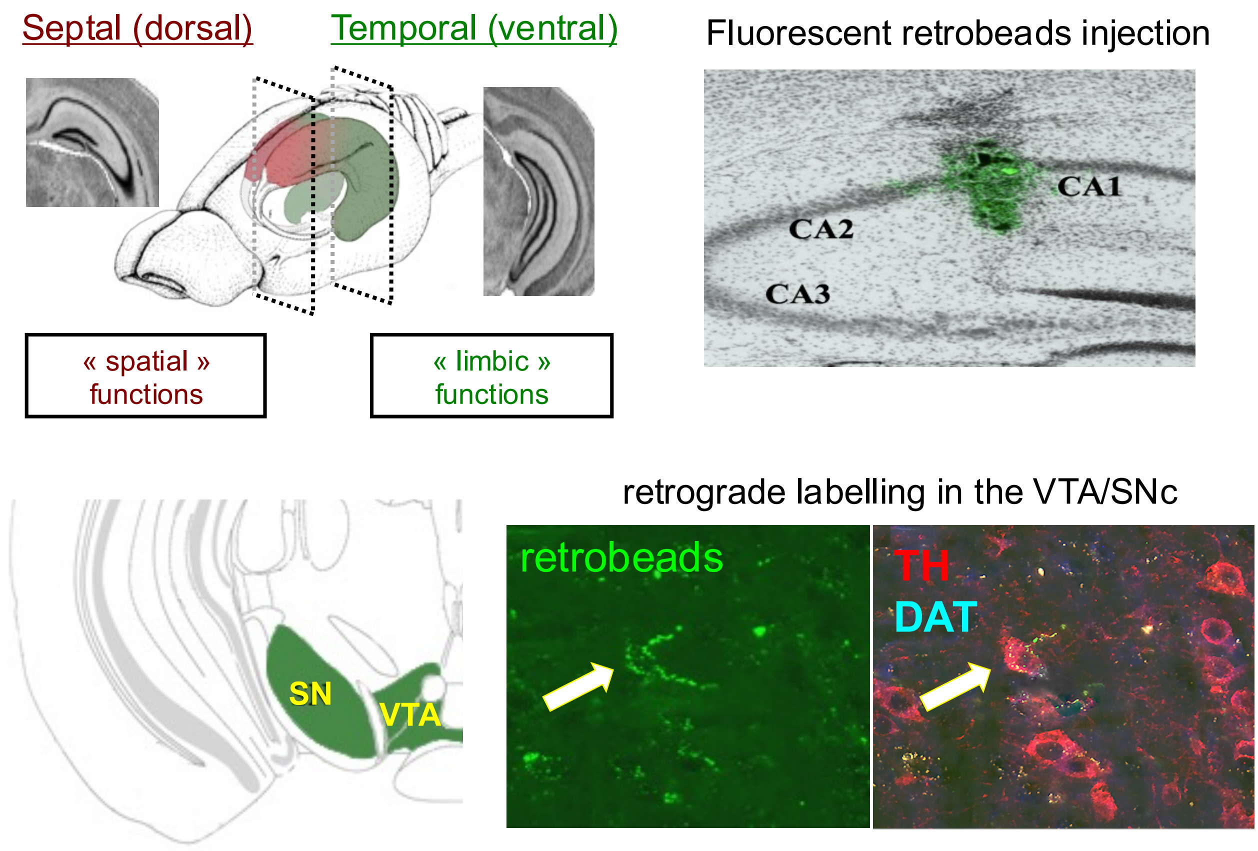


**Figure Sup 1:** Dopamine transporter expression in DA fibers to the Hippocampus.
Top: Schematic 3D representation of mouse HP with septal (upper right) and temporal (bottom right) parts of the HP. Schematic representation of retrobeads injection sites in the temporal HP-CA1. Bottom: Co-immunolabeling for TH (red) and DAT (blue) was carried out to identify if the retrolabeled neurons in the VTA and SNc were positive for TH and/or DAT
DAT, dopamine transporter; HP, hippocampus; TH, tyrosine hydroxylase; VTA, Ventral Tegmental Area; SN, Substancia Nigra;

### Chronic blockade of DAT in the septal hippocampus did not produce cognitive deficits

GBR12935 (300nmol/L) was chronically infused in animals cannulated for the septal part of the hippocampus following the same procedure as for the temporal hippocampus. Cognitive performances of the mice in the spatial MWM, the NORT and the OPRT were assessed.

For the spatial MWM, mice were subjected to 2 training phases; an initial acquisition phase of 7 days with one platform position then reversal training was performed within the next 6 days to assess for behavioral flexibility. A probe test was performed 24 hours after last training. Animals were infused with GBR12935 each day 30 min before the first training session as previously. Both NaCl and GBR-treated mice learned properly during the first acquisition phase and during the reversal phase. Latency to escape decreased similarly in the two groups over the 7 days of initial training phase and 6 following days of reversal training (Figure Sup 2 A,B). After reversal acquisition, GBR-infused mice were exploring the target quadrant significantly more than the controls (NaCl, 33.3± 4 sec; GBR, 42.1±8 sec, what is the P value?). NaCl group, although showing a trend, did not reach full significance in the probe test after reversal ($p=.09$ compared with 25%) but considering the adequate learning in both training phases of NaCl-treated mice, and the successful performance of GBR-treated mice, we decided to not replicate the experiment (Figure Sup 2 C). After 2 weeks of GBR infusion in septal coordinates, both NaCl- and GBR-treated mice displayed successful novelty and spatial recognition memory in the NORT (NaCl, .61±.02; GBR, .65±.06) and the OPRT (NaCl, .66±.04; GBR, .61±.02) respectively (Figure Sup 2 D,E).

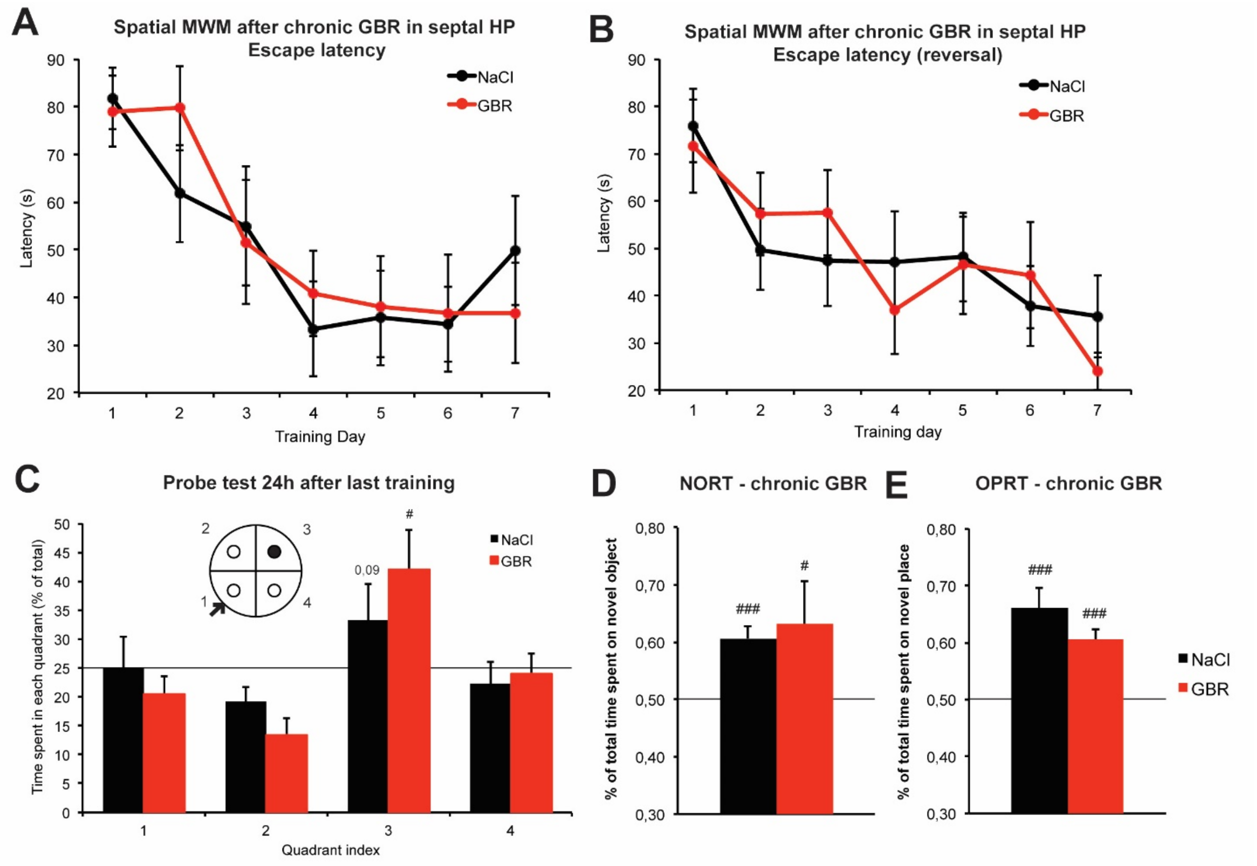


**Figure Sup 2. Effect of GBR12935 intrahippocampal chronic infusion in the septal/dorsal HP on spatial and recognition memory.**

**A)** Escape latency in the MWM with daily infusion of GBR (30 min before first trial, mean escape latencies over two trials a day). After 7 days of training, both NaCl and GBR treated animals properly acquired the platform location (n=10, 9). **B)** Reversal of the platform location. Escape latency in the MWM after the platform was moved from quadrant 1 to quadrant 3. After 7 days of training, both NaCl and GBR treated animals properly acquired the new platform location. **C)** Probe test of spatial memory 24 hours after the last training of the reversal learning phase. NaCl treated animals spent almost significantly more time in the target quadrant (p=.09 compared with 25%). GBR animals spent significantly more time in the target quadrant (#p<.05 compared with 25%). **D)** NORT scores after 2 weeks of chronic GBR infusion in septal HP. Both groups displayed significant visuospatial retention of the older object (n=6 ,6; #p=.05 and ###p<.001 compared with .50; p=.51 between groups). **E)** OPRT scores after 2 weeks of chronic GBR infusion in septal HP. Both groups displayed significant visuospatial retention of the older place (n=6 6; ###p<.001 compared with .50; p=.21 between groups). Bars are SEM.

### Supplemental methods

#### Stereotaxic surgeries

Mice were deeply anaesthetized with isoflurane ($O_2$ debit: 1 mL/min.; isoflurane 5%) then maintained under moderate isoflurane concentration (1-2%) for the entire surgery (about 45 minutes). Anesthetized mice were placed in a stereotaxic apparatus (Stoelting, "Just For Mice" digital stereotaxic apparatus). Scalp skin was cut and removed using cotton tips soaked in iodine solution. Bilateral cannula positioning was marked at appropriate coordinates related to the skull bregma. Holes were drilled through the top skull bone and meninges gently torn with the tip of a syringe. 2 small screws (Morris Co., F00CE125) were placed in a more anterior or posterior location depending of the cannula type to help secure the apparatus in place along further manipulations. Cannulas were gently inserted inside the brain at the appropriate depth. Luting cement was applied on the skull bone to enhance adhesion (C&B-Metabond Quick! Luting cement, Parkell prod.). Dental cement (Jet Denture Repair, Lang Dental) was applied all around the cannula and open space on the skull to maintain it in place. Once cement was dry, animals were placed in a recovery cage with heating pad and moist food. Animals were kept undisturbed for 2 weeks in their home cage and treated with topical antibiotic cream (Neosporin) to prevent infection around the surgery scars and cannulas.

#### Immunohistolabeling and microscopy

For dopamine transporter expression evaluation, animals were anesthetized with isoflurane and their brains was quickly dissected and immerged in freezing isopentane solution. The brains were sliced in 12 µm coronal sections with a cryostat, directly mounted on Superfrost Plus slides (Fisher), and maintained at −80°C until processed. A standard immunolabelling protocol [1] was carried out on all brain slices containing the hippocampus. DA transporter (DAT) antibody (1/4000, Millipore, USA) was applied overnight and detected with an anti-rat coupled to Alexa 555 (1/1000, Invitrogen, USA). Hoescht staining was used to facilitate the detection of hippocampal layers and then the slices were coverslipped with Fluoromount-G. Nonspecific binding was evaluated by using DATKO mice (not shown). Sections containing the hippocampus were observed with an AxioObserver microscope (Zeiss, Germany), and all areas containing DAT positive fibers were imaged at 20x.The images were then analyzed using ImageJ software (NIH, USA) and DAT positive fibers were counted in all subdivisions of the hippocampus.

#### Intrahippocampal infusions

The infusion system was composed of two-syringe infusion/withdrawal pumps (SP210iwZ, WPI), Hamilton 10 µL precision syringes (26Ga) and adequate tubing. The tubing was connected to custom internal cannulas fitting with the bilateral external cannula of the animals to excess the external from .5µm (PlasticOne). For infusions, animals were awake, freely moving in an empty cage box with bedding. Drugs were infused in one HP after the other for 1 min at a .5µL/min rate in each side. For the dual infusion experiment, sulpiride was infused 10 minutes before GBR. Animals were put back to their home cage for 30 minutes prior any behavioral habituation or testing.

#### Pharmacology

For behavioral experiments, $D_2$/$D_3$ antagonist (S)-(-)-Sulpiride (Sigma-Aldrich) was suspended in .1mol/L HCl solution, sonicated, then PH of the solution was adjusted at 7.2 with NaOH at 5mg/mL final concentration. NaCl solution (.9%) was commercially purchased. Its PH was adjusted to 7.2 with HCl and NaOH before use. GBR12935 dihydrocloride (Tocris Bioscience) was suspended in sterile $H_2O$ at a stock concentration of 3µmol/L kept frozen at -80°C. GBR was then diluted daily before infusions at the final concentration of 300nmol/L in NaCl solution (.9%, PH 7.2). Solutions were filtered using 0.22µm filters.

For electrophysiological experiments, sulpiride and GBR were prepared as previously described. Sulpiride was suspended and stocked at a 100mmol/L concentration at -80°C before use. $D_1/D_5$ antagonist R-(+)-SCH23390 hydrochloride (Sigma Aldricht) was suspended at a 10mM concentration and kept at -80°C before use. All drugs were diluted the day of the experiment and bath applied through the perfusion medium.

**Behavior**

*Morris Water Maze (MWM):* The water maze consisted of a circular stainless-steel pool (150 cm diameter, 29 cm height) filled with water maintained at 20–22°C and made opaque with the addition of white aqueous emulsion. A circular escape platform (10 cm diameter) was submerged 1 cm below the water surface. During training phase of the spatial version, mice were trained to find the fixed position of the hidden platform, using extra-maze cues. Mice were released from pseudo-randomly assigned start locations and allowed to swim for up to 90 s. They were manually guided to the platform at the end of the trial in case of failure. Animals received one habituation trial on the first day and then two trials per day during 6 to 10 days (90 min inter-trial interval). On the seventh or eleventh day, the platform was removed, and mice were allowed to swim during 60 s from the opposite starting location. For the reversal procedure, training phase (except habituation) and probe testing were repeated but the platform was moved to the opposite quadrant. For the cued / associative version of the MWM, the pool was surrounded by white curtains to occlude sight of extra-maze cues. The platform was signaled by means of an 8 cm high dark bubble placed onto it. Both platform and mice starting point were pseudo-randomly assigned to different locations across trials. Animals received two trials per day during 5 days (90 min ITI).

*Novel-Object Recognition Task (NORT)*: Mice were tested in an open field (25 cm x 25 cm x 25 cm) with 1 cm sawdust on the floor. The open field contained visible cues of different color and shape on each side. On habituation / training day, mice were allowed to explore two identical objects for 5 min. On the test day, i.e. 24 h later, one of the earlier identical objects was placed into the arena at one location and a new object at the other. mice were allowed to explore the familiar and novel objects for 5 min. Memory of the familiar object is associated with increased exploration of the new object. Exploration score was calculated as a performance ratio: time exploring the novel object/ total time of exploration of both objects. Successful novelty exploration was verified by a ratio significantly superior to the symmetric exploration score (.50). Between two trials, the open field and the objects were cleaned with water containing gentle soap then dried. Sawdust also was discarded and replaced before each trial.

*Object-Place Recognition Task (OPRT)*: 24 hours after NORT, mice were tested in the same open field (25 cm x 25 cm x 25 cm) with 1 cm sawdust on the floor. On habituation / training day, mice were allowed to explore two identical objects for 5 min. On testing day, 24 h later, the first object was placed at the same coordinates than habituation and the other one was placed in a new corner of the open field. The mice were allowed to explore the new object configuration for 5 min. Memory of the familiar coordinates is associated with increased exploration of the object that have been misplaced. Exploration score was calculated as a performance ratio: time exploring the novel place/ total time of exploration of both objects. Successful spatial recognition was verified by a ratio significantly superior to the symmetric exploration score (.50). When the whole NORT and OPRT procedure was repeated two times on the same groups of animals with a 2 weeks delay, pairs of objects were different in each test (Figure 2A). Same pair of objects was never used on same animals but same pairs of objects were used in distinct experiments at similar time points of the infusion procedure (1-4 days vs. 15-18 days of infusion) to allow better comparisons between experiments. Animals exploring less than 10s within 5 minutes of presentation - either in the habituation or the testing phase - were systematically excluded.

**Electrophysiology**

*Slice Preparation:* Adult C57BL6/J mice (3-5 months old) were decapitated under deep isoflurane anesthesia. The brain was quickly removed and placed in a cold slicing solution (concentrations in mmol/L: KCl 2.5, CaCl2 .1, MgCl2 4, KH2PO4 1.25, NaHCO3 26, glucose 10, sucrose 25.2) bubbled with 95% O2/5% CO2 gas mixture. Coronal slices (300 µm) containing the temporal hippocampus were cut and placed on a perfused chamber filled with aCSF (concentrations in mmol/L: NaCl 125, KCl 2.5, CaCl2 2, MgCl2 2, NaH2PO4 1.25, NaHCO3 26, glucose 25). Slices were incubated 90 minutes in a 32°C bath and allowed to recover for 1 hour at room temperature before the experiment.

*Long-Term Potentiation (LTP) / Long-Term Depression (LTD):* Field excitatory postsynaptic potentials (fEPSPs) were obtained from extracellular recordings in the *stratum radiatum* layer of the CA1 region of slices with a glass electrode filled with aCSF (2.7 – 3.3 MΩ) at room temperature. fEPSP was evoked every 20 s by 80 µs test stimulation at afferent Schaffer's collateral fibers. Amplitude of the fEPSPs was recorded for increasing intensities of stimulation to build an input/output curve. Intensity was then adjusted to obtain amplitudes around 50% of the maximum response for LTP and around 70% for LTD. During all the experiment, $GABA_A$ antagonist bicuculline (5µmol/L) was added to the perfusion medium (aCSF). Dopaminergic drugs were diluted in the perfusion medium from stock solutions and bath-applied for 30 to 40 min. prior to stimulation. LTP was induced by a high frequency tetanic stimulation (HFS) consisting of 3 trains of 100 pulses at a frequency of 100 Hz separated by 20 s each. LTD was induced by application of 1200 pairs of pulses with an interpulse interval of 200 ms at a low frequency of 1 Hz (ppLFS). After application of the stimulation protocol, at time 0, drugs were washed-out except for bicuculline and test stimulation was resumed for 60 min. Arrows indicate HFS or ppLFS stimulations.

1 Dal Bo G, St-Gelais F, Danik M, Williams S, Cotton M, Trudeau LE. Dopamine neurons in culture express VGLUT2 explaining their capacity to release glutamate at synapses in addition to dopamine. J Neurochem. 2004;88(6):1398-405.
